# Short report: First whole-genome evidence of dengue virus in field-caught mosquitoes from southern Brazil

**DOI:** 10.1101/2025.10.22.684033

**Authors:** Amanda Cupertino de Freitas, Sara Cândida Ferreira dos Santos, Luiz Marcelo R. Tomé, Vagner Fonseca, Talita Émile Ribeiro Adelino, Natália R. Guimarães, Felipe C.M. Iani, Keldenn Moreno, Bruna Regina Diniz Souza, Getúlio Dornelles Souza, Ellen Caroline Nobre Santos, Lívia Victória Rodrigues Baldon, Rafaela Luiza Moreira, Maria Eduarda Calazans Rodrigues, Isaque João da Silva de Faria, João Paulo Pereira de Almeida, Luiz Carlos Junior Alcantara, Marta Giovanetti, Álvaro Gil Araujo Ferreira

**Affiliations:** Instituto René Rachou, Fundação Oswaldo Cruz, Brazil, 30190-009; Ecovec, Brazil, 31310-260; Programa Interunidades de Pós-graduação em Bioinformática, Universidade Federal de Minas Gerais, Brazil, 31270-901; Fundação Ezequiel Dias, Brazil, 30510-010; Departamento de Ciências Exatas e da Terra, Universidade do Estado da Bahia, Brazil, 41192-010; Vigilância de Roedores e Vetores da Secretaria Municipal de Saúde (CGVS/SMS), Porto Alegre, Brazil, 90810-240; Departamento de Bioquímica e Imunologia, Instituto de Ciências Biológicas, Universidade Federal de Minas Gerais, Brazil, 31270-901; Sciences and Technologies for Sustainable Development and One Health, Università Campus Bio-Medico di Roma, Italy, 00128; Oswaldo Cruz Institute, Oswaldo Cruz Foundation, Rio de Janeiro, Brazil

## Abstract

Dengue virus (DENV) is a major global health threat whose expansion into temperate regions has been facilitated by climate change and vector adaptation. Despite recurrent epidemics in Brazil, genomic surveillance in mosquitoes remains limited, particularly in the South. To address this gap, we implemented a novel urban mosquito-trapping strategy optimized for low-density and peri-domestic environments in Porto Alegre, Rio Grande do Sul. Between April and July 2023, were collected 4,768 *Aedes aegypti* samples across 16 neighborhoods, generating 2,022 pools. Among these, 41 pools tested positive for DENV, of which 33 pools exhibited RNA integrity suitable for sequencing. Whole-genome sequencing revealed 33 DENV-positive pools, including 29 DENV-1 (Genotype V) and 4 DENV-2 (Genotype II). Two pools contained both serotypes, highlighting the risk of sequential infections in humans. Phylogenetic analyses indicated sustained local transmission alongside multiple introductions, while recurrent mutations, particularly in NS1, NS2A, and NS5, suggested ongoing viral adaptation. These findings represent the first vector-based genomic data for dengue in southern Brazil and demonstrate the utility of mosquito genomics for early outbreak detection, serotype monitoring, and preparedness in emerging transmission zones.

**Author Summary:** Dengue virus (DENV) continues to expand into new regions, driven by climate change and mosquito adaptation. In Brazil, dengue epidemics are recurrent, but genomic surveillance of mosquitoes is still scarce, particularly in the southern states. To fill this gap, we implemented an urban mosquito-trapping strategy designed for low-density and peri-domestic environments in Porto Alegre, Rio Grande do Sul. From April to July 2023, we collected 4,768 *Aedes aegypti* samples across 16 neighborhoods, generating 2,022 pools. Among these, 41 pools were positive for DENV, of which 33 pools exhibited RNA integrity suitable for sequencing, mainly DENV-1 (Genotype V) and DENV-2 (Genotype II), with two pools showing co-circulation of both serotypes, raising concern for sequential infections in humans. Phylogenetic analysis revealed sustained local transmission combined with multiple virus introductions, and mutations in NS1, NS2A, and NS5 suggested ongoing viral adaptation. Our findings provide the first mosquito-based genomic data for dengue in southern Brazil and highlight how mosquito genomics can strengthen early outbreak detection, serotype monitoring, and epidemic preparedness.

## Background

Dengue fever is a mosquito-borne disease caused by *Orthoflavivirus denguei* (DENV), a single-stranded, positive-sense RNA virus in the *Flaviviridae* family. It comprises four serotypes (DENV-1 to DENV-4), each with multiple genotypes. Transmission occurs via infected *Aedes* mosquitoes—primarily *Aedes aegypti* and *Aedes albopictus*—and can lead to a wide range of clinical outcomes, from mild febrile illness to severe disease and death [1,5]. Environmental and anthropogenic factors such as climate change, urbanization, deforestation, and human mobility have facilitated the global expansion of dengue. Temperature, in particular, plays a central role in viral transmission and vector competence. Optimal transmission occurs between 32°C and 33°C, although infection can persist at lower thresholds [1,5]. Elevated temperatures can enhance mosquito activity, modify morphology, and improve viral replication by affecting structural proteins such as the envelope (E) protein. Studies have also shown that increasing temperatures improve *Aedes albopictus*’ ability to transmit DENV-2 by facilitating viral dissemination from the midgut to the salivary glands [1,5].

DENV was first reported in Brazil during epidemics in 1846–1848 and 1851–1853, with additional outbreaks in 1916 and 1923 [2,3]. After re-emerging in 1982 in Roraima, major outbreaks followed, notably in Rio de Janeiro in 1986 with the introduction of DENV-1 and the first detection of *Aedes albopictus* in the country [2]. Subsequent introductions included DENV-2 in 1990, DENV-3 in 2000, and DENV-4 in 2012–2013. In 2019, Brazil reported over 1.5 million cases and 782 deaths [3]. While the Southeast, Northeast, and Central-West are hyperendemic, the South—particularly Rio Grande do Sul (RS)—was historically less affected due to its temperate climate. However, climate change and vector adaptation have driven dengue expansion in the region [4]. *Aedes aegypti* was first detected in RS in 1995; the first autochthonous case followed in 2007. The virus has since spread to 453 of 497 municipalities by 2022 (4). From 2021–2022, RS recorded 101,481 cases and 121 deaths [4]. In 2023, 38,176 cases and 54 deaths were confirmed. By 2024, RS experienced its largest outbreak with 207,465 confirmed cases, a 9.2% case-fatality rate, and *Aedes aegypti* found in 94.9% of municipalities [8,9]. Despite this, genomic surveillance remains limited, especially in mosquitoes, hindering our understanding of serotype circulation, transmission, and viral evolution in underrepresented regions such as southern Brazil. To address this gap, we implemented a novel urban mosquito-trapping strategy specifically designed for low-density and peri-domestic environments.

## The study

Between April and July 2023, we collected 4,768 *Aedes aegypti* mosquitoes across 16 neighborhoods of Porto Alegre, spanning five macro-regions defined by the city’s epidemiological planning framework: East Zone (Partenon, Jardim Botânico, Santa Rosa de Lima), North Zone (Sarandi, São Sebastião, Passo das Pedras), South Zone (Tristeza, Nonoai), Central-East Zone (Santa Tereza), and Northeast Zone (Vila João Pessoa, Vila São José, Mário Quintana, Bom Jesus, Costa e Silva, Vila Ipiranga) (**Figure 1a**). Our sampling strategy prioritized ecological and socio-spatial diversity and was optimized to detect vectors in urban zones typically underrepresented in entomological surveillance. Following collection, mosquitoes were grouped into 2,022 pools based on location and collection date. Among these, 41 pools were positive for DENV, of which 33 pools exhibited RNA integrity suitable for sequencing. (**Table S1**). We performed RNA extraction and high-throughput whole-genome sequencing on all samples to assess DENV presence, determine serotype distribution, and characterize viral genetic diversity within local mosquito populations. Adult *Aedes aegypti* were collected using Mosquitrap® devices (Ecovec, Brazil) deployed at 250-meter intervals across urban areas. Traps were checked weekly, and mosquitoes were collected from each location. Captured mosquitoes from each trap were pooled (males and females combined) into tubes containing 250 µL of lysis buffer (8.2 M guanidine thiocyanate, 80 mM Tris-HCl, 70 mM EDTA, pH 8.0) with ten 0.1 mm zirconium beads. Mechanical homogenization was performed using a FastPrep-96™ system (1,800 rpm, 60 s).

**Figure 1.**
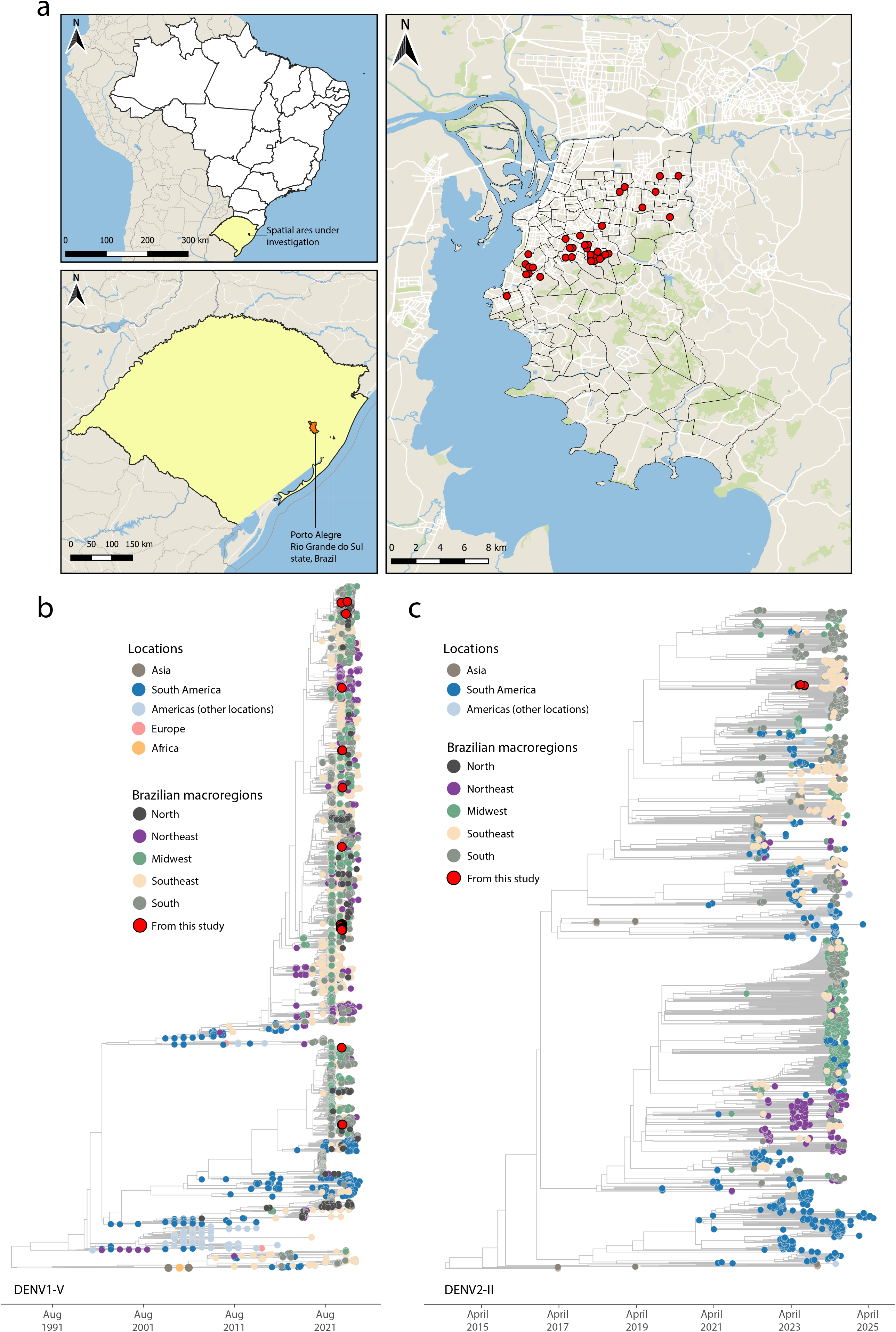
Phylogenetic and spatial analysis of DENV from Aedes mosquitoes in Porto Alegre, Brazil. A) Maps showing the spatial area under investigation. Red dots indicate mosquito collection sites; B) Maximum likelihood (ML) phylogenetic tree of 29 complete DENV-1 genome sequences generated in this study, analyzed alongside 5,635 reference sequences from GISAID and GenBank. Colors indicate sampling locations by global region and Brazilian macroregion; C) ML phylogenetic tree of 4 complete DENV-2 genome sequences generated in this study, analyzed with 1,830 reference sequences from GISAID and GenBank. Colors indicate sampling locations.

Total RNA was extracted from homogenized mosquito samples using the BioGene Viral DNA/RNA Extraction Kit (Bioclin, Brazil), following the manufacturer’s protocol for silica-column purification. The RNA was eluted in 20 µL RNase-free water and was first screened by RT-qPCR to detect DENV serotypes 1–4 and select candidates for for virus genome sequencing [10, 11, 12]. Complementary DNA (cDNA) was synthesized using SuperScript IV Reverse Transcriptase (Thermo Fisher Scientific, Waltham, MA, USA), followed by amplification via 35-cycle multiplex PCR with Q5 High-Fidelity Hot-Start DNA Polymerase (New England Biolabs, Ipswich, MA, USA) and serotype-specific primers targeting full DENV genomes [10, 11]. Amplicons were purified using 1× AMPure XP beads (Beckman Coulter, Brea, CA, USA) and quantified with a Qubit 3.0 fluorometer and Qubit™ dsDNA HS Assay Kit (Thermo Fisher Scientific) [10, 12]. Sequencing libraries were prepared using the Illumina COVIDSeq Test, following the manufacturer’s protocol, and sequenced on the MiSeq platform with a 300-cycle MiSeq Reagent Kit v2. Raw reads were trimmed using Trimmomatic 0.39 and mapped to the reference genome (GISAID: EPI_ISL_17983078 and EPI_ISL_19609373) with Minimap2 2.30-r1287 Subsequent processing included sorting and indexing with Samtools 1.22, indel correction with Pilon 1.24. Low-depth positions (<5×) were masked as ‘N’, and only genomes with >60% coverage were retained for phylogenetic analysis [13]. Sequences were aligned using MAFFT v7.505 and manually inspected in AliView v1.30 to ensure alignment quality [14]. A maximum likelihood tree was inferred using IQ-TREE2 with the GTR+G4+F model. Temporal signal was assessed with TempEst v41, and outliers were removed to improve dating accuracy. The final time-scaled phylogeny was reconstructed with TreeTime using a constant rate model, and a discrete trait analysis was performed to infer viral movements across Brazilian regions and globally [13,14].

Of the 41 mosquito pools positive for DENV, each representing the total number of mosquitoes collected from a single trap over a 7-day period, ranging from 1 to 12 individuals and totaling 149 mosquitoes. Thirty-three yielded sufficient RNA for whole-genome sequencing and tested positive for DENV, 29 for DENV-1 and 4 for DENV-2. This marks the first genomic detection of dengue virus in field-caught mosquitoes from southern Brazil. Two pools (ID_LAB: 4542341, 4539959) showed co-detection of both serotypes, indicating simultaneous circulation of DENV-1 and DENV-2 within the same vector population. While this does not confirm co-infection at the individual mosquito level, it has epidemiological relevance due to the risk of sequential infections in humans. Positive samples showed Ct values ranging from 19.79 to 37, with most below 25, reflecting high viral loads. Sequencing depth ranged from 54× to 2,081×, with read counts between 8,090 and 222,964 for each sequenced sample. Genome coverage ranged from 61.6% to 93.0%, with over 75% of samples achieving >85% coverage, supporting reliable genotyping and phylogenetic analysis (**Table S1**).

All DENV-1 genomes were classified as Genotype V (lineages E.1 and D.1.1) and all DENV-2 genomes as Genotype II (lineage F.1.1.2), both previously reported in Brazil [6,7]. DENV-1 was widely distributed across Porto Alegre’s five macro-regions, while DENV-2 was geographically limited to the East and Northeast zones (**Figure 1a**). Phylogenetic analysis revealed that DENV-1 sequences from this study clustered into four distinct clades (**Figure 1b**). Most grouped with sequences from southern Brazil, consistent with ongoing local transmission, while others clustered with viruses from different regions, suggesting multiple introductions and regional viral exchange. DENV-2 sequences were nested among those previously sampled from the South, preceding the 2024 outbreak across southern and southeastern Brazil (**Figure 1c**).

Additionally, all generated genomes were screened for mutations and annotated using SnpEff to predict potential functional impacts. Most mutations in DENV-1 and DENV-2 genomes were synonymous, with low predicted impact as they do not alter the amino acid sequence of viral proteins. Missense mutations, which can have a moderate impact through amino acid substitutions, were identified in both serotypes. In DENV-1, these were restricted to non-structural proteins—NS1 (n = 4), NS2A (n = 3), NS3 (n = 1), NS4B (n = 1), and NS5 (n = 1)—with NS1 and NS2A most frequently affected (Table S2). In DENV-2, missense mutations occurred in both structural (E, n = 3) and non-structural proteins—NS1 (n = 1), NS2A (n = 1), NS3 (n = 2), NS4B (n = 1), and NS5 (n = 5)—with the highest frequency observed in NS5 (Table S3). Many of these mutations, however, were not unique to our dataset; they have been reported in Brazilian genomes from both current and past epidemics, suggesting a long-standing trend toward viral adaptation. Although their precise functional implications remain to be determined, the recurrent occurrence of changes in proteins involved in replication and immune evasion may reflect selective pressures acting on dengue viruses in Brazil—a hypothesis that warrants further investigation.

These results provide the first vector-based whole-genome data on dengue virus diversity in southern Brazil, offering evidence of probable active transmission of both DENV-1 and DENV-2 in local *Aedes* populations. The high sequencing depth and genome coverage achieved support the detection of co-circulating serotypes, underscoring the need to extend surveillance beyond clinical case reporting, particularly in areas previously considered marginal for dengue transmission. As dengue expands southward, vector genomics emerges as a critical tool for early warning, outbreak mitigation, and understanding urban transmission dynamics under changing climatic and epidemiological conditions. Moreover, genomic surveillance facilitates the identification of mutations with potential roles in immune evasion or increased infectivity, informing timely public health interventions such as vaccine updates and targeted vector control. Collectively, these findings highlight the dynamic and interconnected nature of dengue transmission and the strategic importance of integrating vector-based genomics into epidemic preparedness frameworks.

## Ethical Approval statement

Not Applicable.

## References

1. Branda, F et al. Dengue virus transmission in Italy: historical trends up to 2023 and a data repository into the future. Scientific data, v. 11, n. 1, p. 1–11, 2024.

2. Pinheiro, F. (1997). Re-emergence of dengue and emergence of dengue haemorrhagic fever in the Americas.

3. Adelino, T.É.R., Giovanetti, M., Fonseca, V., Xavier, J., de Abreu, Á.S., do Nascimento, V. A., … & Alcantara, L. C. J. (2021). Field and classroom initiatives for portable sequence-based monitoring of dengue virus in Brazil. Nature communications, 12(1), 2296.

4. Baes Pereira, S., Conrad Bohm, B., dos Reis Gomes, A., Aguiar Gonçalves, G., Campos Pereira Bruhn, N., Silva Belo, V., … & Pascoti Bruhn, F. R. (2025). Emergence and spatiotemporal incidence of dengue in Rio Grande do Sul, Brazil. Scientific Reports, 15(1), 18933.

5. Liu, Z., Zhang, Q., Li, L., He, J., Guo, J., Wang, Z., … & Li, T. (2023). The effect of temperature on dengue virus transmission by Aedes mosquitoes. Frontiers in cellular and infection microbiology, 13, 1242173.

6. Aksamentov, I., Roemer, C., Hodcroft, E. B., & Neher, R. A., (2021). Nextclade: clade assignment, mutation calling and quality control for viral genomes. Journal of Open Source Software, 6(67), 3773, 10.21105/joss.03773

7. Vilsker M, Moosa Y, Nooij S, Fonseca V, Ghysens Y, Dumon K, et al. Genome Detective: an automated system for virus identification from high-throughput sequencing data. Birol I, editor. Bioinformatics. 2019;35(5):871–3. Available from: 10.1093/bioinformat-ics/bty695

8. Centro Estadual de Vigilância em Saúde. (2024). Informativo epidemiológico de arboviroses: ano de 2023 – semanas epidemiológicas 01 a 52. CEVS/SES-RS. Disponível em: https://www.cevs.rs.gov.br/arboviroses-informativo-epidemiologico (Acessado em 29/07/2025).

9. Centro Estadual de Vigilância em Saúde. (2025). Informativo epidemiológico semestral: Dengue e outras arboviroses – SE 01 a 52/2024. CEVS/SES-RS. Disponível em: https://www.cevs.rs.gov.br/arboviroses-informativo-epidemiologico (Acessado em 29/07/2025)

10. Adelino, T.É.R., Giovanetti, M., Fonseca, V., Xavier, J., de Abreu, Á.S., do Nascimento, V. A., … & Alcantara, L. C. J. (2021). Field and classroom initiatives for portable sequence-based monitoring of dengue virus in Brazil. Nature communications, 12(1), 2296.

11. Johnson BW, Russell BJ, Lanciotti RS. Serotype-specific detection of dengue viruses in a fourplex real-time reverse transcriptase PCR assay. J Clin Microbiol. 2005 Oct;43(10):4977–83. doi: 10.1128/JCM.43.10.4977-4983.2005. PMID: 16207951; PMCID: PMC1248506.

12. Castilho de Arruda, L. D., Giovanetti, M., Fonseca, V., Zardin, M. C. S. U., Lichs, G. G. D. C., Asato, S., … & Cavalheiro Maymone Gonçalves, C. (2023). Dengue fever surveillance in Mato Grosso do Sul: insights from genomic analysis and implications for public health strategies. Viruses, 15(9), 1790.

13. Giovanetti, M., Slavov, S. N., Fonseca, V., Wilkinson, E., Tegally, H., Patané, J. S. L., … & Covas, D.T. (2022). Genomic epidemiology of the SARS-CoV-2 epidemic in Brazil. Nature Microbiology, 7(9), 1490–1500.

14. Giovanetti, M., Micheli, V., Mancon, A., Mileto, D., & Rizzo, A. (2025). Phylogenetic analysis of Chandipura virus: insights from a preliminary genomic study. International Journal of Molecular Sciences, 26(3), 1021.

15. Li, Q., & Kang, C. (2022). Structures and dynamics of dengue virus nonstructural membrane proteins. Membranes, 12(2), 231.

